# Hierarchical Brain Network for Face and Voice Integration of Emotion Expression

**DOI:** 10.1101/197426

**Authors:** Jodie Davies-Thompson, Giulia V. Elli, Mohamed Rezk, Stefania Benetti, Markus van Ackeren, Olivier Collignon

## Abstract

The brain has separate specialized computational units to process faces and voices located in occipital and temporal cortices. However, humans seamlessly integrate signals from the faces and voices of others for optimal social interaction. How are emotional expressions, when delivered by different sensory modalities (faces and voices), integrated in the brain? In this study, we characterized the brains’ response to faces, voices, and combined face-voice information (congruent, incongruent), which varied in expression (neutral, fearful). Using a whole-brain approach, we found that only the right posterior superior temporal sulcus (rpSTS) responded more to bimodal stimuli than to face or voice alone but only when the stimuli contained emotional expression. Face-and voice-selective regions of interest extracted from independent functional localizers, similarly revealed multisensory integration in the face-selective rpSTS only; further, this was the only face-selective region that also responded significantly to voices. Dynamic Causal Modeling revealed that the rpSTS receives unidirectional information from the face-selective fusiform face area (FFA), and voice-selective temporal voice area (TVA), with emotional expression affecting the connection strength. Our study promotes a hierarchical model of face and voice integration, with convergence in the rpSTS, and that such integration depends on the (emotional) salience of the stimuli.

## INTRODUCTION

The ability to quickly and accurately recognize emotional expressions is a fundamental cognitive skill for effective social interactions. In particular, the tone of the voice and the facial expression of the communicator are two crucial cues that we use to appropriately orient our behavior in a social context. Typically, emotion is simultaneously expressed through separate sensory channels, like the face and the voice, and this information is seamlessly integrated in a coherent percept producing enhanced discrimination and faster reaction times (Massaro and Egan 1996; De Gelder and Vroomen 2000; Collignon et al. 2008). Further, perceiving a facial expression can alter the percept of a vocal emotional expression (Pye and Bestelmeyer 2015), and emotional input from one unattended sensory input can systematically influence the judgment of the attended sensory stream (De Gelder and Vroomen 2000; Collignon *et al.* 2008). Additionally, audiovisual integration of emotion expressions can arise pre-attentively (Föcker et al. 2011) and is unconstrained by available attentional resources (Vroomen et al. 2001). Together, this suggests that the integration of facial and vocal emotion information may be a rather automatic process (Vroomen *et al.* 2001). However, the neural mechanisms underlying how emotional signals from the face and the voice are integrated, remains unclear. More specifically, it is debated whether integration of facial and vocal emotion information occurs within a distinct convergence zone (Kreifelts et al. 2009; Watson, Latinus, Noguchi, et al. 2014) or could already be observed within the face and voice-selective networks (Blank et al. 2011; Joassin et al. 2011).

Influential models of identity recognition suggest two largely distinct brain networks for face and voice processing, with a shared multisensory convergence zone for the integration of information across sensory modalities (Bruce and Young 1986; Burton et al. 1990; Haxby et al. 2000; Watson, Latinus, Charest, et al. 2014). Some studies have suggested that the integration of facial and vocal emotional signals occur in multisensory regions like the amygdala, orbitofrontal cortex, and posterior superior temporal sulcus (STS) (Romanski 2007; Kreifelts *et al.* 2009; Peelen et al. 2010; Watson, Latinus, Charest, *et al.* 2014; Hölig et al. 2017); regions separate from the core face and voice selective network (Campanella and Belin 2007). However, the absence of independent localisers in these studies makes it difficult to ascertain whether regions showing multisensory integration are located within face- or voice-selective regions, or instead occurs within an independent convergence zone. It is therefore crucial to localise face- and voice-selective regions independently, and examine the response to visual, auditory, and visuo-auditory emotion information, in order to determine whether multisensory integration of face and voice signals occur in independent converging zones or can already be observed in face- or voice-selective regions.

Alternatively, integration of emotional information of faces and voices could occur at early stages of processing via reciprocal connections between regions across the face and voice preferential networks. Recent models of identity recognition have indeed suggested that integration of faces and voices could occur within unisensory regions (Von Kriegstein et al. 2005; von Kriegstein and Giraud 2006; Blank *et al.* 2011; Joassin *et al.* 2011) as a result of direct structural (Blank *et al.* 2011) and functional (Von Kriegstein *et al.* 2005) connectivity between face-selective regions in the fusiform cortex (fusiform face area, FFA; (Kanwisher et al. 1997) and voice-selective regions in the middle temporal gyrus (temporal voice area, TVA; (Belin et al. 2000; Belin et al. 2002). Interestingly, electrode studies in macaques have suggested that a significant proportion of neurons in voice-selective regions in auditory cortex respond more to audiovisual stimuli as compared to unimodal vocal or facial sitmuli (Ghazanfar et al. 2005; Perrodin et al. 2014), and are sensitive to asynchronies in the onset of faces and voices (Perrodin et al. 2015).

Finally, face and voice regions may together form a general social processing network, with all regions involved in the processing of both visual and auditory human stimuli. For example, viewing point-light displays depicting biological motion or geometric shapes depicting social interactions appear to activate both the face-selective FFA and posterior STS (Bonda et al. 1996; Castelli et al. 2000; Grossman and Blake 2002; Schultz et al. 2003) suggesting that these regions may play a more abstract role in social perception. Such a model would propose that so-called ‘unisensory’ regions would not only respond to their preferred stimulus, but also respond to both unisensory facial and vocal expressions of emotion.

Our study was specifically designed to test these contrasting views on whether integration of information from faces and voices occurs in earlier or later stages of processing, and how emotional expressions alter this integration. First, using two independent face and voice localisers, we assessed whether ‘unisensory’ regions respond to input from different sensory modalities, by examining the response to faces within the voice network and to voices within the face network. Second, the same subjects underwent an additional scan to examine the response to neutral and fearful unimodal faces, unimodal voices, and bimodal face and voice stimuli. The goal of this second experiment was to test whether the functionally selective face and voice regions defined with the localizers are involved in the multisensory integration of emotional stimuli. Third, we examined the response to unimodal and bimodal stimuli in a whole brain analysis in order to determine the existence of a distinct multisensory convergence zone. Finally, we use dynamic causal modeling (DCM) to examine how face and voice information flows and is integrated across these regions.

## MATERIALS AND METHODS

### Participants

24 healthy right-handed participants (12 females; mean age: 26 years, SD: 5) with no history of neurological dysfunction, and with self-reported normal or corrected-to-normal vision, and normal hearing took part in the study. The protocol was approved by the institutional review boards of the University of Trento and the Center for Mind/Brain Sciences (CIMeC), and written informed consent was obtained for all subjects in accordance with The Code of Ethics of the World Medical Association, Declaration of Helsinki (Rickham, 1964).

### Imaging parameters

Subjects were scanned in a Bruker BioSpin MedSpec 4T scanner at the Center of Mind/Brain Sciences (CIMeC), University of Trento. T2*-weighted scans using echo planar imaging were used to collect data for all functional scans from 37 sequential axial slices (TR = 2200ms, TE = 30ms, FA = 76°, FOV = 192mm, 3mm slice thickness, voxel size = 3×3mm). These were co-registered onto a T1-weighted 3D MPRAGE anatomical image (TR = 2700ms, TE = 4.18ms, 176 axial slices, FOV = 256mm, 1mm slice thickness, voxel size = 1×1mm), from each participant.

### Procedure

Participants undertook two visits. During the first visit, participants undertook a 90-minute scan session during which they completed a face localizer, voice localizer, and the multisensory experiment in the same fixed order. Identical scans were conducted during the second visit.

### Face localizer

To identify regions responding preferentially to faces, a face localiser scan was carried out for each subject. Subjects viewed blocks containing images from one of the four different categories: neutral faces, fearful faces, Fourier scrambled faces, or objects. Face images were 20 front-on photographs of different actors (10 female) with either neutral or fearful expressions, taken from the Radboud Faces Database (Langner et al. 2010). Faces were cropped to their natural chin and hairline in order to remove external features. The object stimuli consisted of 20 pictures of objects (e.g., cars, houses, etc.). The scrambled face stimuli were created by applying a Fourier phase randomization procedure to the neutral face images, which preserves only the low-level properties (e.g., luminance, spatial frequency, colors) (e.g.(Sadr and Sinha 2004)). All images were framed in a white rectangle (220×270 pixels) and had comparable mean luminance and contrast (measured using SHINE Toolbox in MATLAB – (Willenbockel et al. 2010). In order to minimize an after-image effect, visual stimuli were presented centrally but with slight changes in their position around the fixation point, varying either in the x- or in the y-axis by 30 pixels. Each image was presented for 1s followed by a 50ms black screen. There were 20 images per block with each block lasting approximately 21s. Each condition was repeated 8 times. Blocks were separated by a 7-9s fixation blank screen (mean = 8s; jitter = 1s) and were presented in a fixed order (neutral face, object, fearful face, scrambled faces). The localizer was split into two runs: each lasting approximately 8 minutes. Subjects performed a one-back task, pressing a button when the same stimulus was repeated twice in a row.

### Voice localizer

To identify regions responding preferentially to voices, a voice localiser scan was carried out for each subject. Subjects were presented with blocks containing sounds from one of the five different categories: neutral voices, fearful voices, scrambled neutral voices, scrambled fearful voices, or objects sounds. The voice stimuli consisted in the articulation of the vowel /*a*/ recorded by 20 different actors (10 female) expressing either neutral or fearful emotion (taken from the Montreal Affective Voices – (Belin et al. 2008)). The object sound stimuli were non-verbal object sounds referring to non-living objects, namely human action sounds (e.g. lighting a match), bells and musical instruments (e.g. Christmas bells) and automated machinery (e.g. hair dryer).

Scrambled versions of the vocal and object sounds were obtained using MATLAB (The MathWorks, Inc., Natick, Massachusetts, United States). Scrambling was inspired by the method of Belin and colleagues (2000,2002) but differed in that the scrambling of amplitude and phase components was conducted separately within frequency windows instead of time windows (see (Dormal et al. 2017). Each vocal and object sound was submitted to a fast Fourier transformation and the resulting components were separated into frequency windows of ∼700 Hz based on their center frequency. Scrambling was then performed by randomly intermixing the magnitude and phase of each Fourier component (Belin *et al.* 2000; Belin *et al.* 2002) within each of these frequency windows separately. The inverse Fourier transform was then applied on the resulting signal. The output was a sound of the same length of the original sound with similar energy within each frequency band. For scrambled vocal sounds only, the envelope of the original voice was further applied on the output signal. This was not done for scrambled object sounds because the application of the original envelope in this case led to recognition of many scrambled object sounds despite the scrambling. Hence, for these sounds, a 5ms ramp was applied in the beginning and at the end and a 5ms silence was added at the beginning. Following standard practice, voices, object sounds and their scrambled versions were equalized in root mean square (RMS) level.

Each auditory stimulus lasted for 1s followed by 50ms of silence. There were 20 auditory stimuli per block with each block lasting approximately 21s. Each condition was repeated 8 times. Blocks were separated by a 7-9s fixation blank screen (mean = 8s; jitter = 1s) and were presented in a fixed order (neutral voice, scrambled neutral voice, fearful voice, scrambled fearful voice, object sounds). The localizer was split into two runs, each lasting approximately 9 minutes. Subjects were asked to respond via button press when two sequentially presented stimuli were identical.

### Multisensory experiment

There were eight experimental conditions which consisted of 4 unimodal conditions: (i) *neutral faces (FaceN)*; (ii) *fearful faces (FaceF)*; (iii) *neutral voices (VoiceN)*; (iv) *fearful voices (VoiceF)*; and 4 bimodal conditions: (v) congruent neutral faces with neutral voices (*CongN*); (vi) congruent fearful faces with fearful voices (*CongF*); (vii) incongruent neutral faces with fearful voices (*IncongNF-FV*); (viii) incongruent fearful faces with neutral voices (*IncongFF-NV*). Figure 1 shows examples of each condition.

**Figure 1.**
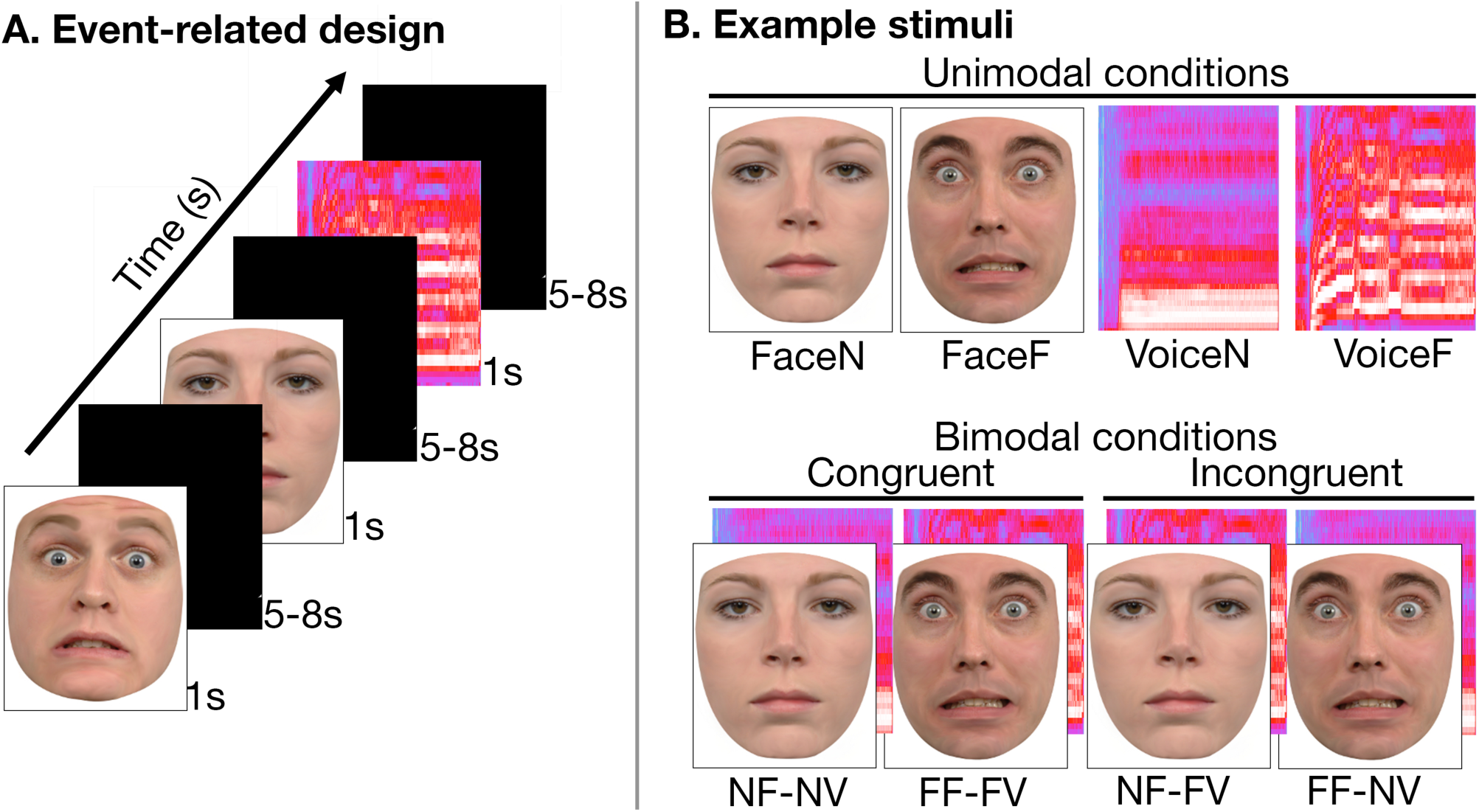
Stimuli presentation and example stimuli used in the experiment. Stimuli were presented in an event-related design. Each run lasted ∼15 minutes, and were repeated 3 times. Face (F) and voice (V) stimuli could be either fearful (F) or neutral (N) emotional expressions. In the bimodal condition, voices and faces were presented together as either congruent (neutral face and voice, or fearful face and voice) or incongruent (neutral face and fearful voice, or fearful face and neutral voice).

Stimuli were 28 video clips of 14 professional actors (7 female) pronouncing the vowel /*a*/ while performing either neutral or fearful facial expressions and voice intonations (Simon et al. 2008). The videos were selected from a set of 62 clips based on the ratings by 20 independent judges’ (half of them female) who decided which emotion was performed (fear/neutral) and how representative it was (0 Not at all – 10 Very much). All selected videos were correctly recognized by at least 19 judges, and had a minimum representative mean score of 7. The audio tracks from these videos were used in the unimodal voice conditions, and the videos without the audio tracks were used in the unimodal face conditions. Congruent conditions contained the original video clips, with synced faces and voices. Incongruent conditions were created using the video and audio tracks of opposite emotional expressions performed by the same actor, with facial motion and auditory onset synced. All stimuli (112) were the same duration (1s). All audio signals were equalized in energy using Root Mean Square normalization.

An event-related design was used to present the stimuli. Each trial consisted of a single stimulus from one of the eight conditions presented for 1s, followed by a black screen for 5-8s (mean = 6.5s; jitter = 1s). Each of the 8 condition were repeated 14 times, resulting in a total of 112 trials in a single run lasting approximately 15 minutes. Each run was repeated 3 times, with each of the 3 runs containing stimuli performed by different actors. To monitor attentional load across stimulus conditions, subjects performed an orthogonal gender discrimination task for each stimuli in which they responded via a button press (right middle and index finger), as to whether the stimuli was that of a male or female.

### fMRI Analysis

Analysis of the MRI data was carried out using Statistical Parametric Mapping (SPM8 – Welcome Department of Imaging Neuroscience, 2009), implemented in MATLAB R2008a (The MathWorks, Inc.). The initial 4 functional volumes of data from each scan were removed to minimize the effects of magnetic saturation. Preprocessing of the functional MR data included, in the following order: slice time correction to the middle slice; realignment of functional time series; co-registration with the anatomical data; segmentation of the structural; normalization of the structural and of the functional images into standardized Montreal Neurological Institute (MNI) space; and smoothing (Gaussian kernel, 4mm FWHM) of the functional images.

In the fMRI data analysis, single subjects were entered into a fixed-effect analysis (FFX). Changes in the regional blood oxygenation level dependent (BOLD) signal were estimated using a general linear model (GLM). Regressors of interest consisted of experimental events time-locked to the stimulus onset and convolved with a standard hemodynamic response function. Motion parameters obtained during preprocessing were included as regressors of no interest. A high-pass filter with a cut-off frequency of 128Hz was used to correct for low-frequency noise and signal drift.

### Whole brain analysis (multisensory experiment)

Statistical images from the single-subject level analysis were spatially smoothed (Gaussian kernel 6mm FWHM) and entered into a mixed-effects group-level (RFX) analysis. Statistical inferences were then performed at a threshold of p < 0.05 FWE-corrected for multiple comparisons.

Multisensory regions across the entire brain were defined as responding more to bimodal stimulation as compared to both unimodal conditions, using a conjunction (AND) analysis (bimodal – visual) ∩ (bimodal – auditory) (Van Atteveldt et al. 2004; Ethofer et al. 2013). We analysed the two emotions separately for neutral (*CongN – FaceN*) ∩ (*CongN– VoiceN*), and fearful (*CongF–FaceF*) ∩ (*CongF –VoiceF*) conditions. We chose this criteria, because, unlike the conservative ‘additive model’ (FV > [F+V]) (Perrault et al. 2003, 2005; Stanford et al. 2005) the ‘max criterion’ model ([FV > F] ∩ [FV > V]) better accommodates for negative responses to conditions, whilst also providing a stricter criteria than the more liberal ‘mean criterion’ model (AV > mean(A,V])(Calvert 2001; Beauchamp 2005) in which a greater response to one of the unimodal condition can still result in a region being labeled as multisensory.

To examine the effects of congruency across the brain, we collapsed both congruent conditions (*CongN, CongF*) and compared the response to the collapsed incongruent conditions (*IncongNF-FV, IncongFF-NV).* Note that in this case, it is not straightforward to examine the effects of emotional expression on congruency, as the incongruent conditions cannot be categorized as fearful or neutral.

### Region of interest analysis

To examine the response to the experimental conditions in regions selective for faces and voice, we obtained regions of interest (ROI) at the single-subject level from independent localiser sessions. Face-selective regions of interest (ROI) were identified from the face localizer by the conjunction (*neutral faces* > *scrambled faces*) ∩ (*neutral faces* > *objects*), thresholded at P < 0.001 uncorrected to identify regions responding more to faces than to objects as well as removing low-level image differences. We identified the bilateral fusiform face area (FFA; left: identified in n = 23 participants; right: n = 24), bilateral occipital face area (OFA; left: n = 19; right: n = 20), and the posterior superior temporal sulcus (pSTS; left: n = 11; right: n = 20) in individual subjects (Table 1). These regions are consistent with coordinates from previous studies (Davies-Thompson and Andrews 2012; Rossion et al. 2012). Voice-selective ROIs were identified from the voice localizer by the conjunction (*neutral voices* > *scrambled voices*) ∩ (*neutral voices* > *object sounds*), thresholded at P < 0.001 uncorrected, to identify regions responding more to voices than objects sounds as well as low-level auditory properties. We identified the bilateral temporal voice area (TVA) in middle temporal gyrus (left: n = 16; right: n = 19), a region in the bilateral anterior temporal lobe (ATL; left: n = 12; right: n = 8), and another region in left anterior superior temporal sulcus (aSTS; n = 14) in individual subjects (Table 1). These regions are consistent with reported coordinates from previous studies (Belin *et al.* 2000; Belin *et al.* 2002), While some studies have also identified voice-selective responses in posterior portions of STS (i.e. (Pernet et al. 2015), this region was only observed in a limited number of participants, and was not observed at the group level (see below) – therefore, it was not possible to obtain sufficient data from this region.

**Table 1.**
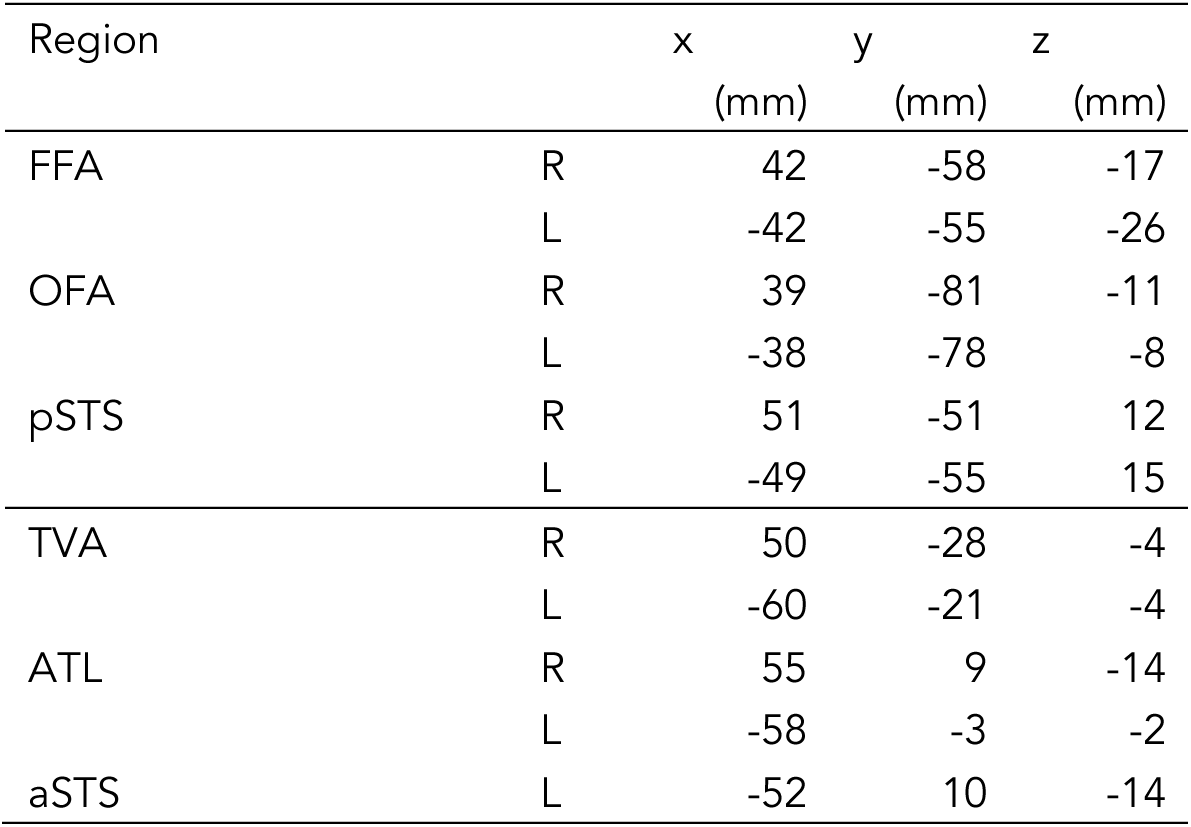
Average MNI coordinates of the face- and voice-selective region of interest analysis.

Given the strict criteria used to identify these regions, some subjects did not have a complete set of regions of interest. In order to obtain data from these regions, which may be subthreshold, we created a 10mm sphere mask around the peak coordinate at the group level and extracted data from individual subjects who did not show this region at the individual subject level. This allowed us to obtain data from all 24 subjects for all regions of interest (Table 1). (Note that analysis showed the same pattern of response for each ROI, regardless of whether regions were defined only at the individual subject level (partial sample) or whether regions were defined from the group level coordinates (whole sample)). The beta values to each of the experiment conditions within the face- and voice-selective regions of interest were then entered into repeated-measures ANOVA to determine significant differences in the response to each stimulus condition.

### Dynamic Causal Modeling

We use Dynamic Causal modeling (DCM; (Friston et al. 2003) to investigate cortical interaction between face and voice-selective regions, and how this gives rise to multisensory responses. In DCM changes in the neuronal states of a set of regions in a network are modeled using a bilinear state equation and a biophysically validated model of how these neuronal dynamics affect the bold response observed in fMRI. The bilinear state equation consists of three main parameters. These parameters are the endogenous interactions between regions (A), modulation of these connections by a stimulus or task (B), and the exogenous input to the network (C).

DCM analysis was implemented using SPM8 toolbox (Wellcome Department of Imaging Neuroscience, London, UK), running in Matlab R2012b (The Math Works, Natick, MA, USA). We defined bilinear models with mean-centered inputs with endogenous connections (A) between the regions of interest. Unimodal faces, unimodal voices, and bimodal face-voice stimuli served as the driving inputs to the regions (C). To investigate how these intrinsic dynamics interact with facial expression, emotional expression (neutral, fearful) was entered as a modulator of connectivity (B) between regions, as well as a direct input (C).

Next, as we were interested in how the intrinsic connections are modulated as a function of emotional expression, each model was repeated for alternative points at which emotional expression might influence the intrinsic connections. In the case of models with bi-directional intrinsic connections (A), modulations (B) were assumed to be bidirectional as well.

To decide which model best explained our results, we used random-effects Bayesian model selection (BMS; (Stephan et al. 2009)).

### Statistical analysis

For each face-selective region (OFA, FFA, pSTS), beta values from the face localiser from individual subjects were entered as the outcome variable in a repeated-measures ANOVA with Hemisphere (left, right), and Condition (neutral faces, emotive faces, objects, scrambled neutral faces) as factors, and Subject as a random effect. To explore the basis of significant main effects and interactions revealed by the ANOVA, two-tailed paired-samples t-tests were then used for post-hoc comparisons (Bonferroni corrected for multiple comparisons). The same method was used to test for differences in the response to the voice localiser within the face-selective regions, with Hemisphere (left, right) and Condition (neutral voices, emotive voices, object sounds, scrambled neutral voices, scrambled emotive voices) as factors, and Subject as a random effect. Identical statistical analyses were conducted for voice-selective regions (TVA, ALT, aSTS), but without Hemisphere as a factor for left aSTS due to no right hemisphere equivalent.

For the multisensory experiment, beta values from face-selective and voice-selective regions were entered as the outcome variable in a repeated-measures ANOVA with Hemisphere (left, right), Emotion (neutral, fearful), and Modality (bimodal congruent, unimodal faces, unimodal voices) as factors, and Subject as a random effect. To explore the basis of interactions and main effects, two-tailed paired-samples t-tests were then used for planned comparisons to compare responses across Modality within each region (Bonferroni corrected for multiple comparisons). Finally, to test for congruency effects in face-selective and voice-selective regions, beta values were entered into independent repeated-measures ANOVAs, with Region (face-selective: 6 regions; voice-selective: 5 regions) and Congruency (congruent, incongruent) as factors, and Subject as a random effect, followed by paired-samples t-tests (Bonferroni corrected).

Finally, to further evaluate the presence or absence of multisensory responses in the face- and voice-selective regions, we conducted Bayesian paired samples t-tests (JASP, Version 0.9) to compare the beta values in the Congruent (bimodal) condition as compared to the each regions preferred stimulus modality (unimodal faces in FFA, OFA, and STS; unimodal voices in MTG and ATL) for both emotion conditions (neutral, fearful). Bayesian statistics have several advantages over classical frequentist approach, notably in being less affected by the problems associated with multiple comparisons (Gelman et al. 2012). Moreover, Bayesian statistic provides evidence, not only in favor of the rejection of the null-hypothesis (presence of evidence for multisensory integration), but also meaningful inferences in favor of the null-hypothesis (absence of evidence for multisensory integration) (Masson 2011; Wagenmakers et al. 2018).

## RESULTS

### Crossmodal selectivity in functional localisers

To determine whether face-selective regions responded to voices, and voice-selective regions responded to faces, we extracted the beta values to all stimuli from the face and voice localisers in all face and voice regions of interest. To specifically test for crossmodal selectivity, beta values to the non-preferred stimuli – the response to the face localiser in voice selective regions, and the response to the voice localiser in face selective regions - were entered into repeated-measures ANOVAs (hemisphere, condition), followed by post-hoc paired-samples t-tests to determine whether: (1) the response to faces was greater than to scrambled faces and objects in voice-selective regions, and (2) the response to voices was greater than scrambled voices and object sounds in face-selective regions.

Figure 2 shows the response to the face and voice conditions in face-selective regions of interest. Independent beta values extracted from the voice localizer experiment only revealed positive beta values (higher than baseline) in the pSTS regions, while other face selective regions (FFA and OFA) showed a pattern of deactivation for auditory categories. A 2 (hemisphere) x 5 (conditions) ANOVA for the voice localiser in the FFA revealed a significant effect of Condition (F(4,92) = 7.24, P < 0.001) but no effect of Hemisphere (F(1,23) = 3.99, P = 0.06) or Interaction (F(4,92) = 2.14, P = 0.08). In both the left and the right FFA, the effect of condition was caused by a negative response to objects as compared to both neutral voices (left: t(23) = 2.33, p < 0.05; right: t(23) = 3.00, p < 0.01) and emotive voices (left: t(23) = 4.24, p < 0.001; right: t(23) = 4.84, p < 0.001); there was also a greater negative response to scrambled emotive voices than to emotive voices in the left FFA (t(23) = 2.32, P < 0.05). The same pattern was observed in the OFA: there was an effect of Condition (F(4,92) = 17.79, P < 0.001) but no effect of Hemisphere (F(1,23) = 0.80, P = 0.38) or Interaction (F(4,92) = 1.20, P = 0.32). This was caused by a negative response to object sounds as compared to both neutral voices (left: t(23) = 4.99, p < 0.001; right: t(23) = 2.65, p < 0.05) and emotive voices (left: t(23) = 7.36, p < 0.001; right: t(23) = 4.89, p < 0.001). In sum, neither the FFA nor OFA responded significantly to human voices. In the pSTS, there was an effect of Condition (F(4,92) = 13.85, p < 0.001) and an Interaction (F(4,92) = 6.15, p < 0.001), but no effect of Hemisphere (F(1,23) = 2.22, p = 0.15). In contrast to the FFA and OFA, the pSTS showed a positive response to both neutral voices > scrambled neutral voices (left: t(23) = 2.90, p < 0.01; right: t(23) = 4.93, p < 0.001) and emotive voices > scrambled emotive voices (left: t(23) = 2.72, p < 0.05; right: t(23) = 4.28, p < 0.001). The right pSTS also responded more to emotive voices > object sounds (t(23) = 2.59, p < 0.05). These data suggest cross-modal selectivity to human voices in the left and right pSTS only.

**Figure 2.**
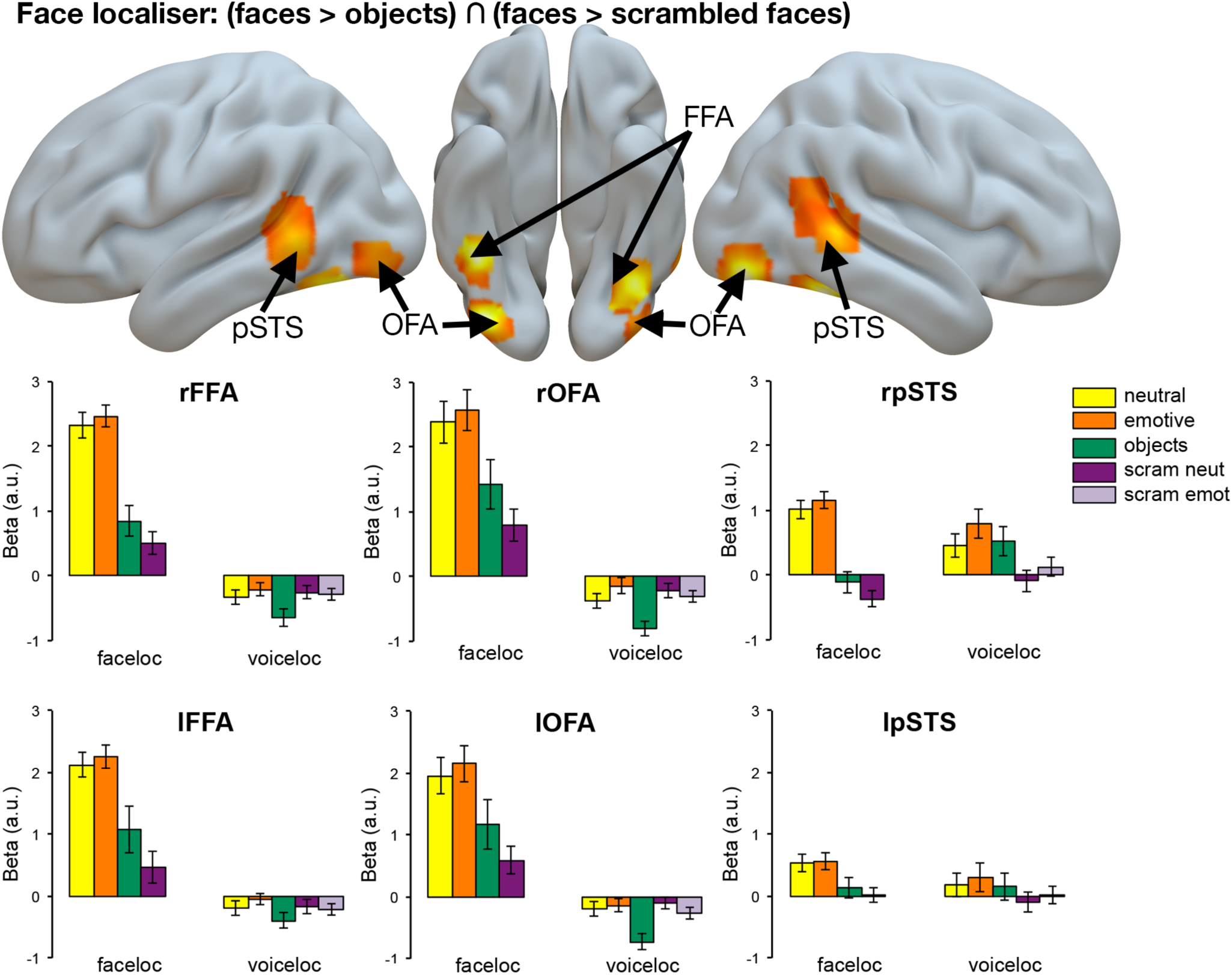
Face-responsive regions identified in the face localizer. Top: Bilateral fusiform face area (FFA), occipital face area (OFA), and a region in the posterior superior temporal sulcus (pSTS) responded more to faces than to scrambled faces and objects. Contrast: (faces > scrambled faces) ∩ (faces > objects). Bottom: The response of these face-selective regions to stimuli presented in the face and voice localisers. Neither the left nor right FFA nor OFA responded significantly to neutral or emotive voices. In contrast, both left and right pSTS responded more to neutral and emotive voices as compared to their scrambled counterparts (neutral voices > scrambled neutral voices; emotive voices > scrambled emotive voices).

Figure 3 shows the response to the face and voice conditions in voice-selective regions of interest. A 2 (hemisphere) × 4 (conditions) ANOVA for the face localiser in the TVA revealed no effect of Condition (F(3,69) = 0.40, p = 0.76) or Hemisphere (F(1,23) = 1.93, p = 0.18), but a significant Interaction (F(3,69) = 3.02, p < 0.05). However, there were no significant differences in the response to neutral or emotive faces as compared to scrambled faces and objects (P’s > 0.17). In ATL, there was no effect of Condition (F(3,69) = 2.03, p = 0.12), Hemisphere (F(1,23) = 0.08, p = 0.79), or Interaction (F(3,69) = 0.34, p = 0.80). Finally, a 1 × 5 (conditions) ANOVA for the left aSTS showed no effect of Condition (F(3,69) = 0.40, p = 0.75). These data suggest no cross-modal selectivity to faces in voice-selective regions.

**Figure 3.**
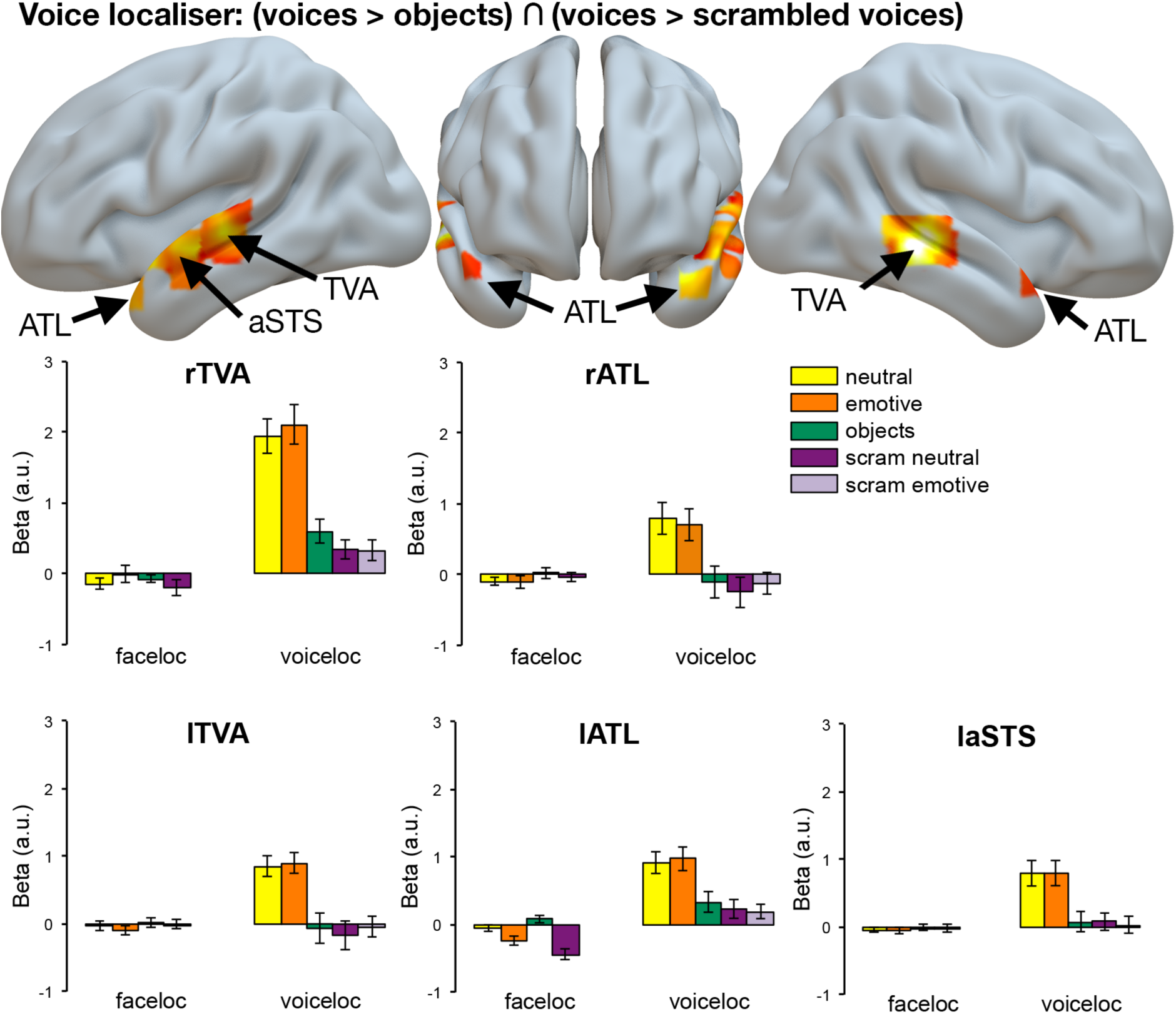
Voice-responsive regions identified in the voice localizer. Top: Bilateral temporal voice area (TVA), anterior temporal lobe (ATL), and a region in the anterior superior temporal sulcus (aSTS) responded more to voices than to scrambled voices and objects. Contrast: (voices > scrambled voices) ∩ (voices > object sounds). Bottom: The response of these voice-selective regions to stimuli presented in the face and voice localisers. Neither the left nor right TVA, ATL, nor left aSTS, responded significantly to neutral or emotive faces.

### Effects of bimodal versus unimodal stimuli

#### Whole brain analysis

Figure 4 (red) shows a region in the right posterior superior temporal sulcus (rpSTS) that responded more to bimodal conditions as compared to unimodal conditions for fearful stimuli in the whole brain analysis (P < 0.05, FWE-corrected for multiple comparisons). However, this region showed no difference in response to bimodal as compared to unimodal neutral stimuli, thus providing evidence of multisensory integration for emotive (fearful) stimuli, but not for neutral stimuli. This region (peak MNI coordinates: 54, −46, 8) overlapped with the face-selective pSTS identified in the independent localizer (Figure 8), and is similar in location to other studies that have suggested a role of this region in multisensory integration (i.e. (Watson, Latinus, Noguchi, *et al.* 2014): 48, −40, 13 and 66, −46, 4; (Kreifelts *et al.* 2009): 54, −42, 10). No other regions responded significantly more to bimodal than to either unimodal conditions for either fearful or neutral stimuli.

**Figure 4.**
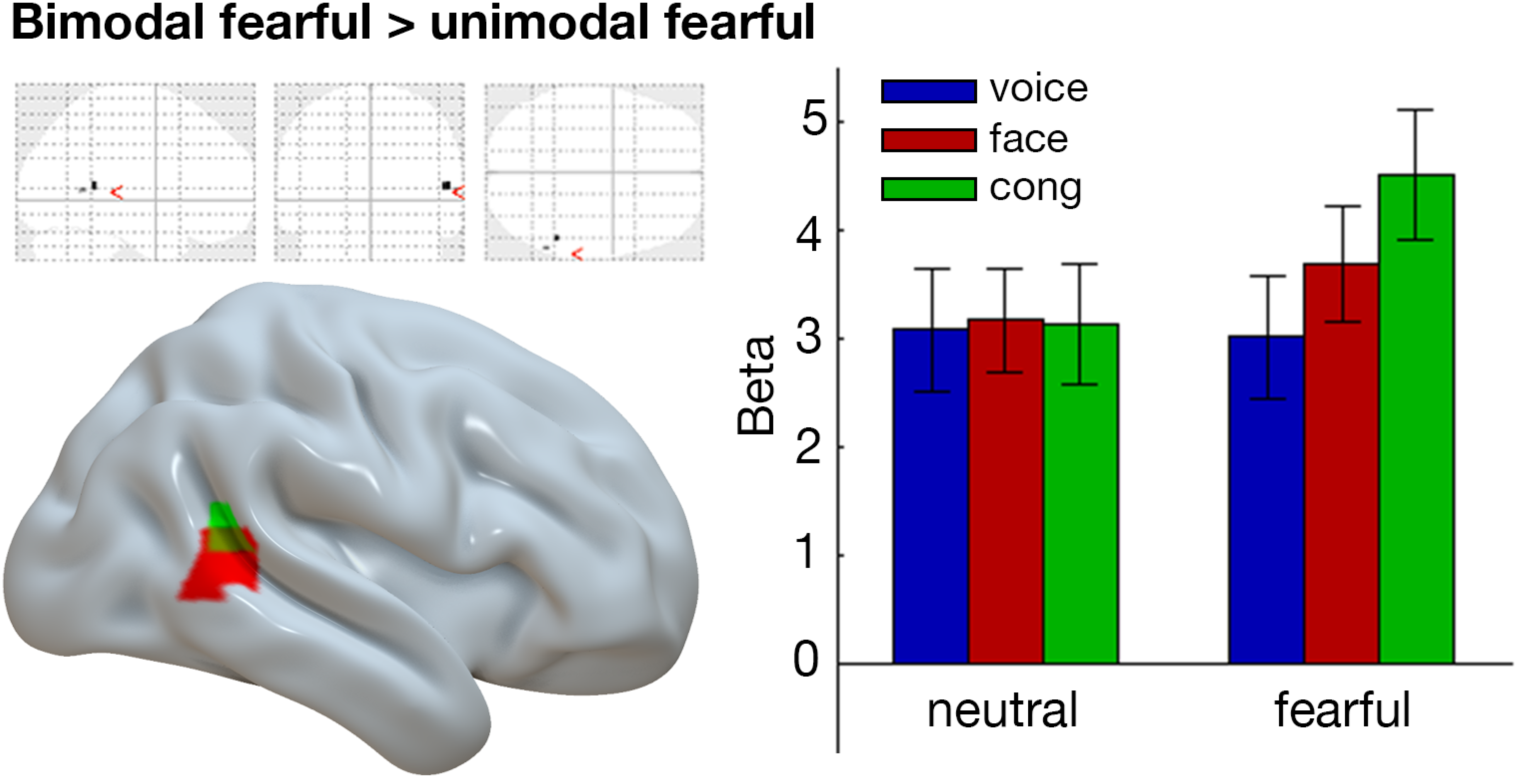
Whole brain analysis. A region in the right superior temporal sulcus (rpSTS; red) responded more to bimodal than to unimodal fearful stimuli. Contrast: [bimodal congruent fear > unimodal fearful face] ∩ [bimodal congruent fear > unimodal fearful voice]. Similarly, a region in the right STS responded significantly more to fear(bimodal > unimodal) > neutral(bimodal > unimodal)(green). Contrast: [(CongF–FaceF) ∩ (CongF – VoiceF)] > [(CongN – FaceN) ∩ (CongN– VoiceN)]. Whole-brain maps are displayed at P < 0.05 FWE-corrected for multiple comparisons.

To further examine the interaction between bimodal to unimodal conditions for fearful versus neutral stimuli, we conducted an additional analysis. Specifically, we first analysed the response to bimodal > unimodal stimuli for the two emotions separately for fearful (*CongF–FaceF*) ∩ (*CongF – VoiceF*) and neutral (*CongN – FaceN*) ∩ (*CongN– VoiceN*) conditions at the individual subject level. Next, these conjunction maps were entered into a paired-samples t-test at the RFX level to specifically test for regions showing an interaction between bimodal > unimodal stimuli across the two emotions. Figure 4 (green) shows one region in the right STS which responded significantly more to fear(bimodal > unimodal) > neutral(bimodal > unimodal) in the whole brain analysis (P < 0.05, FWE-corrected for multiple comparisons). This region partially overlapped with the rSTS region that responded more to bimodal conditions as compared to unimodal conditions for fearful stimuli, providing further support for a role of this region in the multisensory integration of fearful but not neutral face and voice information (see Figure 4).

#### Face-selective regions

Figure 5 shows the response across subjects in face-selective regions of interest (as defined by the conjunction (neutral faces > scrambled faces) ∩ (neutral faces > objects)) to the different conditions in the independent multisensory experiment.

**Figure 5.**
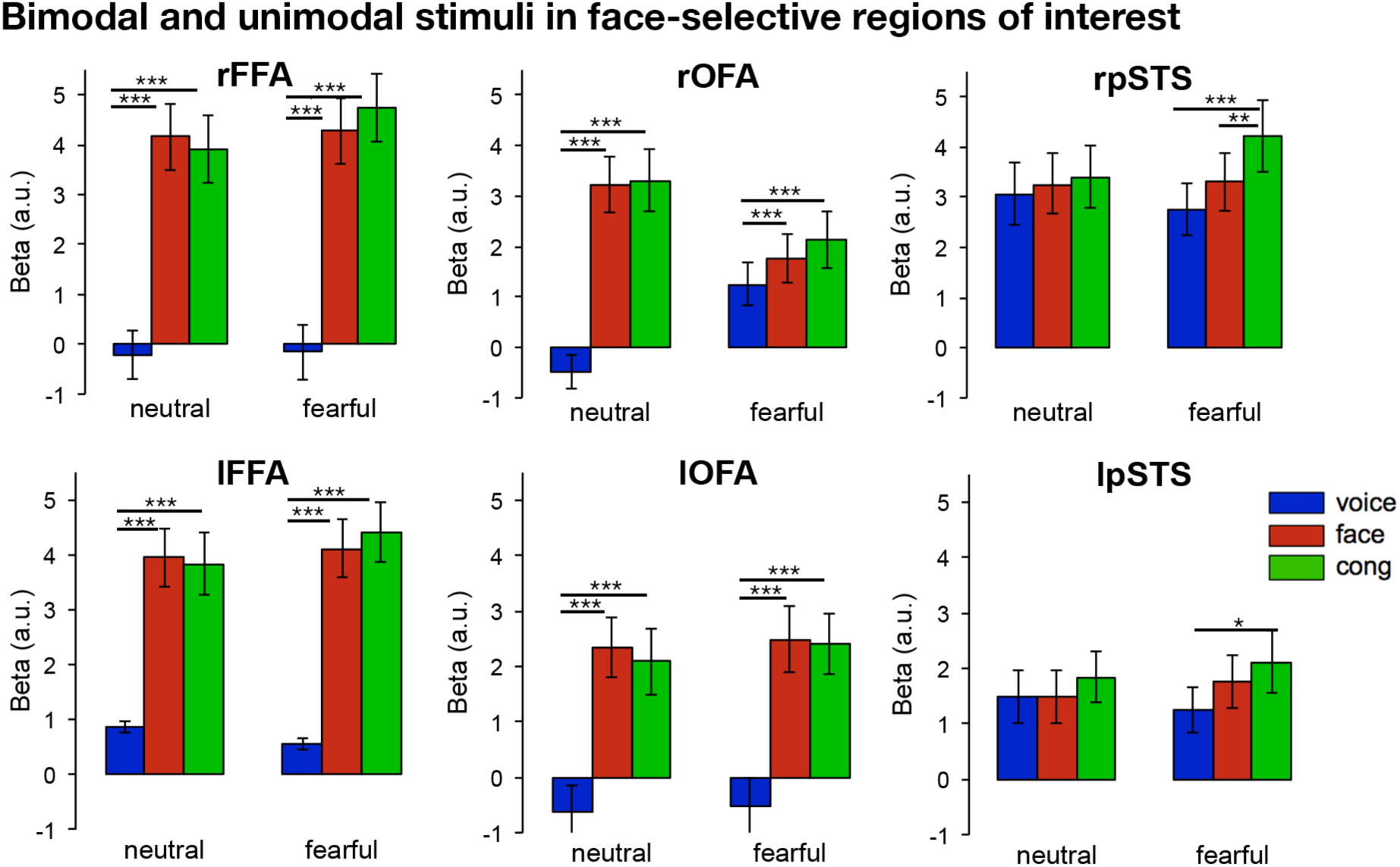
The response to unimodal faces, unimodal voices, and bimodal faces and voices in face-selective regions of interest. The right pSTS responded significantly more to bimodal stimuli (cong) than to unimodal faces and unimodal voices in the fearful conditions. No other face-selective regions responded more to bimodal > unimodal stimuli. *P < 0.05, **P < 0.01, ***P < 0.001.

For both the OFA and FFA, 2 (Hemisphere) × 2 (Emotion: neutral, fearful) x 3 (Modality: face, voice, bimodal) ANOVAs showed effects of Emotion (P’s < 0.05) and Modality (P’s < 0.001), but no effects of Hemisphere (P > 0.10). For all four regions (left FFA, right FFA, left OFA, right OFA), the effect of Modality was due to significantly greater responses to faces and congruent bimodal stimuli as compared to voices for neutral (*neutral faces* > *neutral voices*; *CongN* > *neutral voices*) and fearful stimuli (*fearful faces* > *fearful voices*; *CongF* > *fearful voices*) (P’s < 0.001), but no difference in the response to unimodal faces and congruent bimodal stimuli (*CongN* = *neutral faces*; *CongF* = *fearful faces*) (P’s > 0.07). The effect of Emotion was due to a greater response to *bimodal fearful > bimodal neutral* stimuli in the right hemisphere of the OFA (P = 0.014), and in both the left and right hemisphere of the FFA (P’s < 0.01). Both regions showed a significant interaction between Hemisphere × Modality (P’s < 0.05); in the OFA, this was caused by a greater response in the right > left OFA to bimodal neutral and fearful stimuli (P’s < 0.05); in the FFA, this was caused by a greater response to neutral voices in the left > right hemisphere (P < 0.05).

For the pSTS, a 2 × 2 × 3 ANOVA showed an effect of Hemisphere (F(1,23) = 15.52, P < 0.001), and Modality (F(2,46) = 6.13, P < 0.005), but no effect of Emotion (F(1,23) = 1.55, P = 0.23). However, there was an interaction between Emotion x Modality (F(2,46) = 3.81, P < 0.05) and Hemisphere x Emotion x Modality (F(2,46) = 3.71, P < 0.05). These interactions were caused by differences between the response to bimodal stimuli across emotions and hemispheres. For fearful stimuli, the right pSTS responded significantly more to bimodal congruent stimuli than to both unimodal faces (*CongF* > *fearful faces*: t(23) = 3.39, p < 0.005) and unimodal voices (*CongF* > *fearful voices*: t(23) = 3.61, p < 0.001), but no difference in the response between fearful faces and fearful voices (t(23) = 1.45, p = 0.16). Contrarily, for neutral stimuli, there was no difference in the response between any conditions (P’s > 0.24). Further, there was a significantly greater response to bimodal fearful > bimodal neutral stimuli (t(23) = 3.42, p < 0.005). The differences in the response to bimodal as compared to unimodal stimuli as a function of emotion, was further confirmed by the presence of an interaction between Condition and Modality (F(2,46) = 5.60, P < 0.01). Thus, the right pSTS showed multisensory integration for emotive (fearful) stimuli, but not for neutral stimuli. In the left hemisphere, the pSTS showed a significantly greater response to congruent fearful stimuli as compared to fearful voices (*CongF* > *fearful voices*: t(23) = 2.74, p < 0.05). However, there were no other significant differences for either fearful conditions (P ‘s > 0.11) or neutral stimuli conditions (P’s > 0.13).

To further evaluate the presence or absence of multisensory responses in these regions, we compared the response to congruent bimodal stimuli to unimodal voices (the regions preferred stimulus modality) for both emotion conditions (CongN > neutral faces; CongF > fearful faces). For bilateral FFA, OFA, and left STS, the Bayesian Factors varied between 0.22 and 1.06, supporting the absence of evidence and in favor of the null hypothesis (no difference between the response to congruent bimodal stimuli and unimodal faces). The exception to this was for fearful stimuli in the rSTS where we found further evidence for a difference in the response between bimodal and unimodal fearful stimuli (CongF > fearful faces; Bayesian Factor: 15.84).

#### Voice-selective regions

Figure 6 shows the response across subjects in voice-selective regions of interest (as defined by the conjunction (voices > scrambled voices) ∩ (voices > object sounds)) to the different conditions in the independent multisensory experiment.

**Figure 6.**
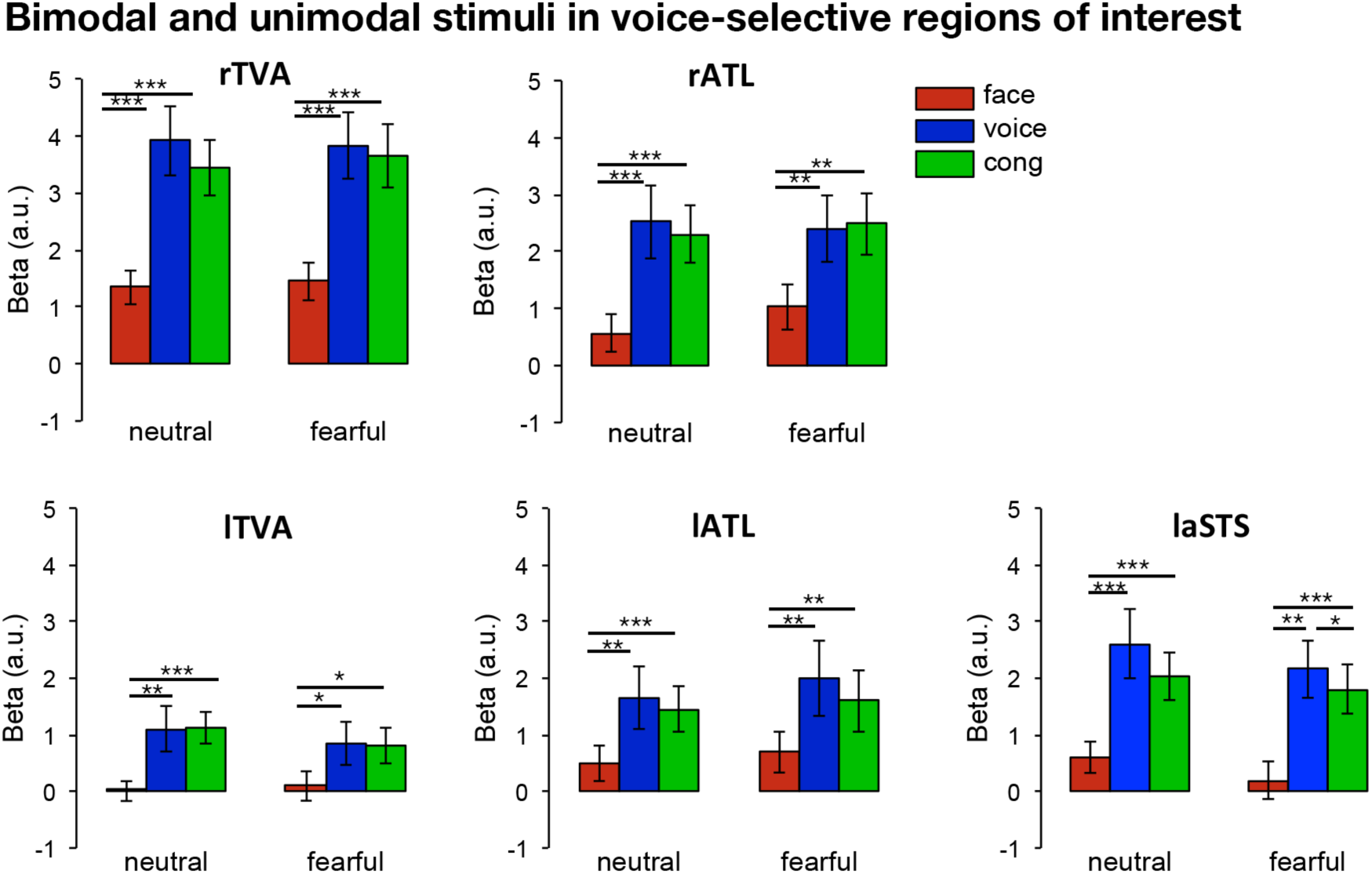
The response to unimodal faces, unimodal voices, and bimodal faces and voices in voice-selective regions of interest. No regions responded more to bimodal (cong) > unimodal faces and unimodal voices. *P < 0.05, **P < 0.01, ***P < 0.001.

2 (Hemisphere) × 2 (Emotion: neutral, fearful) × 3 (Modality: face, voice, bimodal) ANOVAs for the TVA and ATL, showed main effects of Modality (P’s < 0.001), but no effects of Emotion (P’s > 0.71). For all four regions (left TVA, right TVA, left ATL, right ATL), the effect of Modality was due to significantly greater responses to voices and congruent bimodal stimuli as compared to faces for neutral (*neutral voices* > *neutral faces*; *CongN* > *neutral faces*) and fearful stimuli (*fearful voices* > *fearful faces*; *CongF* > *fearful faces*) (P’s < 0.05), but no difference in the response to unimodal voices and congruent bimodal stimuli (*CongN* = *neutral voices*; *CongF* = *fearful voices*) (P’s > 0.18). In TVA, there was also a significant effect of Hemisphere (F(1,23) = 51.95, P < 0.001) and an interaction between Hemisphere x Modality (F(2,46) = 14.63, P < 0.001). This was caused by a greater response in the right > left TVA for all corresponding conditions (i.e. neutral voices in right TVA > neutral voices in left TVA) (P’s < 0.001). No such differences were observed in ATL.

A 2 × 3 ANOVA for the left aSTS showed an effect of Emotion (F(1,23) = 6.82, P < 0.05) and Modality (F(2,46) = 21.63, P < 0.001), but no interaction between Emotion x Modality (F(2,46) = 0.27, P = 0.76). For neutral stimuli, there was a greater response to voices and congruent bimodal stimuli as compared to faces (P’s < 0.001), with no difference in the response between voices and congruent bimodal stimuli (*CongN* = *neutral voices*: t(23) = 1.91, p = 0.07). For fearful stimuli, there was a greater response to voices and congruent bimodal stimuli than to faces (P’s < 0.001); however, there was a significantly greater response to fearful voices as compared to congruent bimodal stimuli (*fearful voices > CongF*: t(23) = 2.14, p < 0.05), showing that this region responds more to fearful voices when presented in isolation as compared to when they were presented with congruent facial stimuli.

To further evaluate the presence or absence of multisensory responses in these regions, we compared the response to congruent bimodal stimuli to unimodal voices (the regions preferred stimulus modality) for both emotion conditions (CongN > neutral voices; CongF > fearful voices). For all 5 regions (bilateral MTG, bilateral ATL, left aSTS), the Bayesian Factors varied between 0.24 and 1.12, supporting the absence of evidence and in favor of the null hypothesis (no difference between the response to congruent bimodal stimuli and unimodal voices).

### Congruency effects

To identify regions that are involved in processing multisensory stimuli as a function of congruency, we examined the response to *congruent > incongruent* stimuli [*(congruent neutral + congruent fearful) – (both incongruent conditions)]* as well as the reverse contrast to show regions responding more to *incongruent > congruent* stimuli [(*both incongruent conditions*) – (*congruent neutral* + *congruent fearful*)].

Figure 7A shows several regions that responded more to *incongruent > congruent* stimuli in a whole brain analysis at a liberal statistical threshold (P < 0.05, FWE-corrected for multiple comparisons). These include regions in bilateral middle frontal gyrus (lMFG, rMFG), bilateral superior parietal cortex (lSupPar, rSupPar), the left middle temporal gyrus, a portion of the left frontal pole (lFront Pole), and a portion of the right pSTS (Table 2). Figure 8 shows that this portion of rpSTS (green) partially overlaps with the face-selective rpSTS (yellow), and to a much lesser extent, the area in rpSTS showing a multimodal effect for fearful stimuli in the group analysis (red). However, with the exception of a trend in rSTS, none of these regions responded significantly more to *bimodal > unimodal* stimuli. In contrast, no regions responded more to congruent than to incongruent stimuli – even at the more liberal (P < 0.001, uncorrected) threshold.

**Figure 7.**
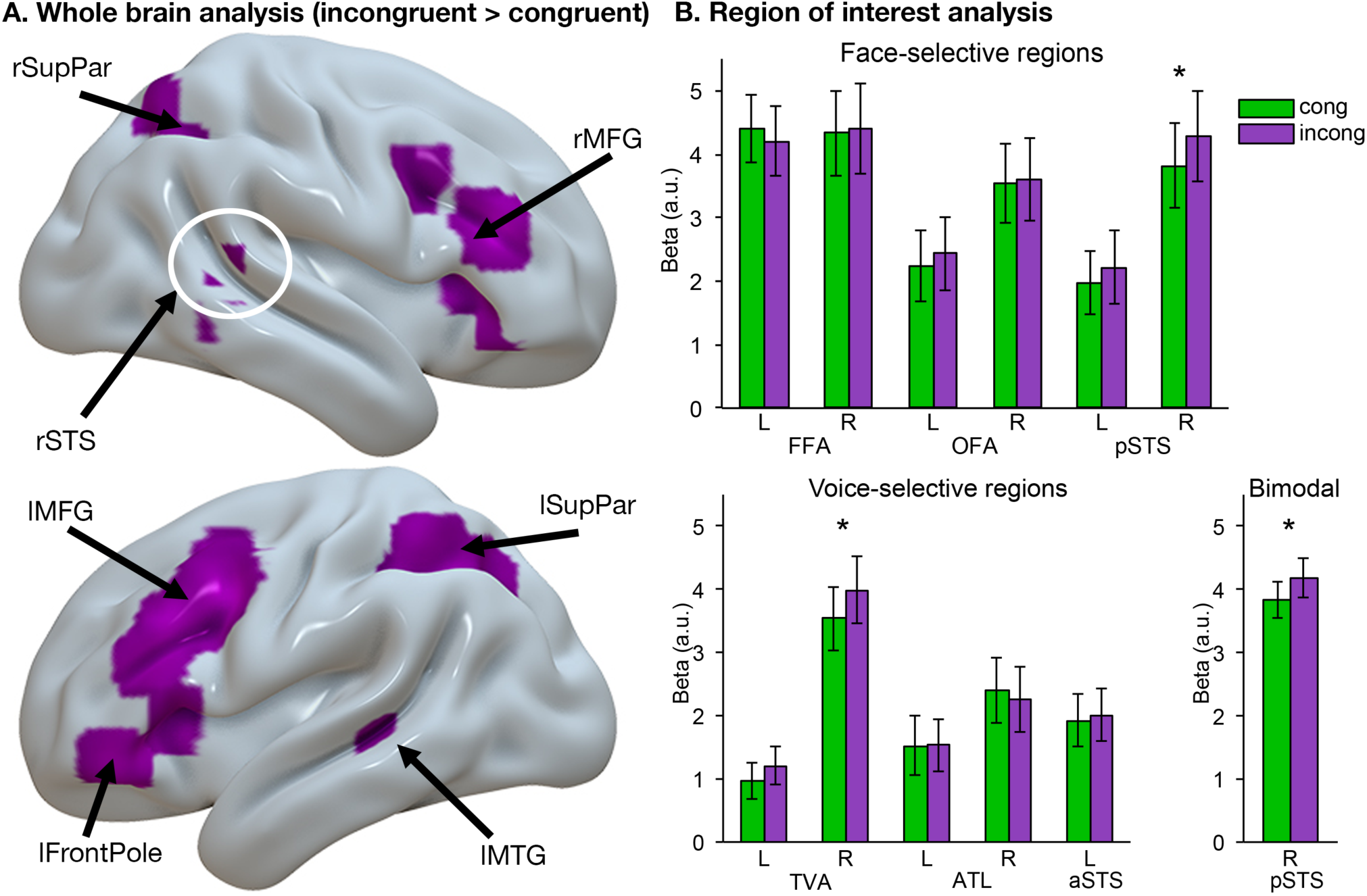
Congruency effects. A). Regions showing congruency effects in the whole brain analysis. Several regions responded more to incongruent > congruent stimuli, including bilateral middle frontal gyrus (lMFG, rMFG), bilateral superior parietal cortex (lSupPar, rSupPar), a portion of the left frontal pole (lFrontPole), and a portion of the rpSTS (indicated by a white circle). Aside from a trend in rpSTS, none of these regions responded more to bimodal > unimodal stimuli. No regions responded more to congruent > incongruent stimuli. Statistical maps thresholded at P < 0.05 FWE-corrected for multiple comparisons. * P < 0.05. B) The response to congruent and incongruent stimuli in the face-selective and voice-selective regions of interest, as well as the region in the group analysis which showed a greater response to bimodal > unimodal stimuli (Figure 4). There was a greater response to incongruent > congruent stimuli in the voice-selective right TVA, the face-selective rpSTS, and the region that responded more to bimodal > unimodal stimuli in the group analysis (Figure 4).

**Table 2.**
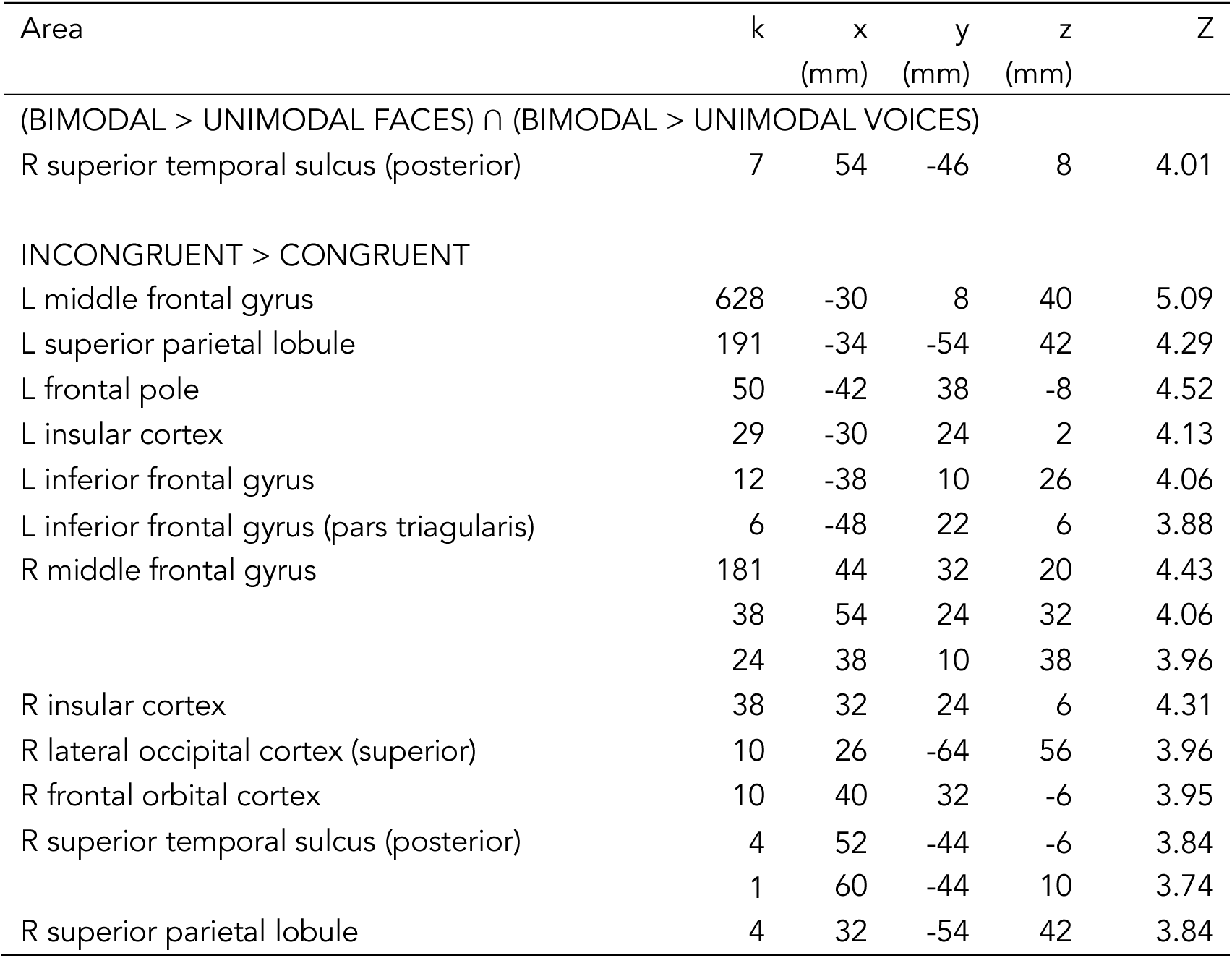
Regions showing congruency effects in the whole brain analysis (P < 0.05, FWE-corrected).

**Table 3.** Bayesian statistics (BF_10_) comparing the response to congruent > unimodal faces in face-selective regions (FFA, OFA, STS), and congruent > unimodal voices in voice-selective regions (MTG, ATL, aSTS).

| BF<sub>10</sub> | Left Hemisphere |  | Right Hemisphere |  |
| --- | --- | --- | --- | --- |
|  | Neutral | Fearful | Neutral | Fearful |
| FFA | .258 | .906 | 1.058 | .332 |
| OFA | .397 | .230 | .224 | .255 |
| STS | .573 | .731 | .241 | **15.836** |
| MTG | .329 | 1.115 | .275 | .240 |
| ATL | .329 | 1.115 | .275 | .240 |
| aSTS | .760 | .876 |  |  |

**Figure 8.**
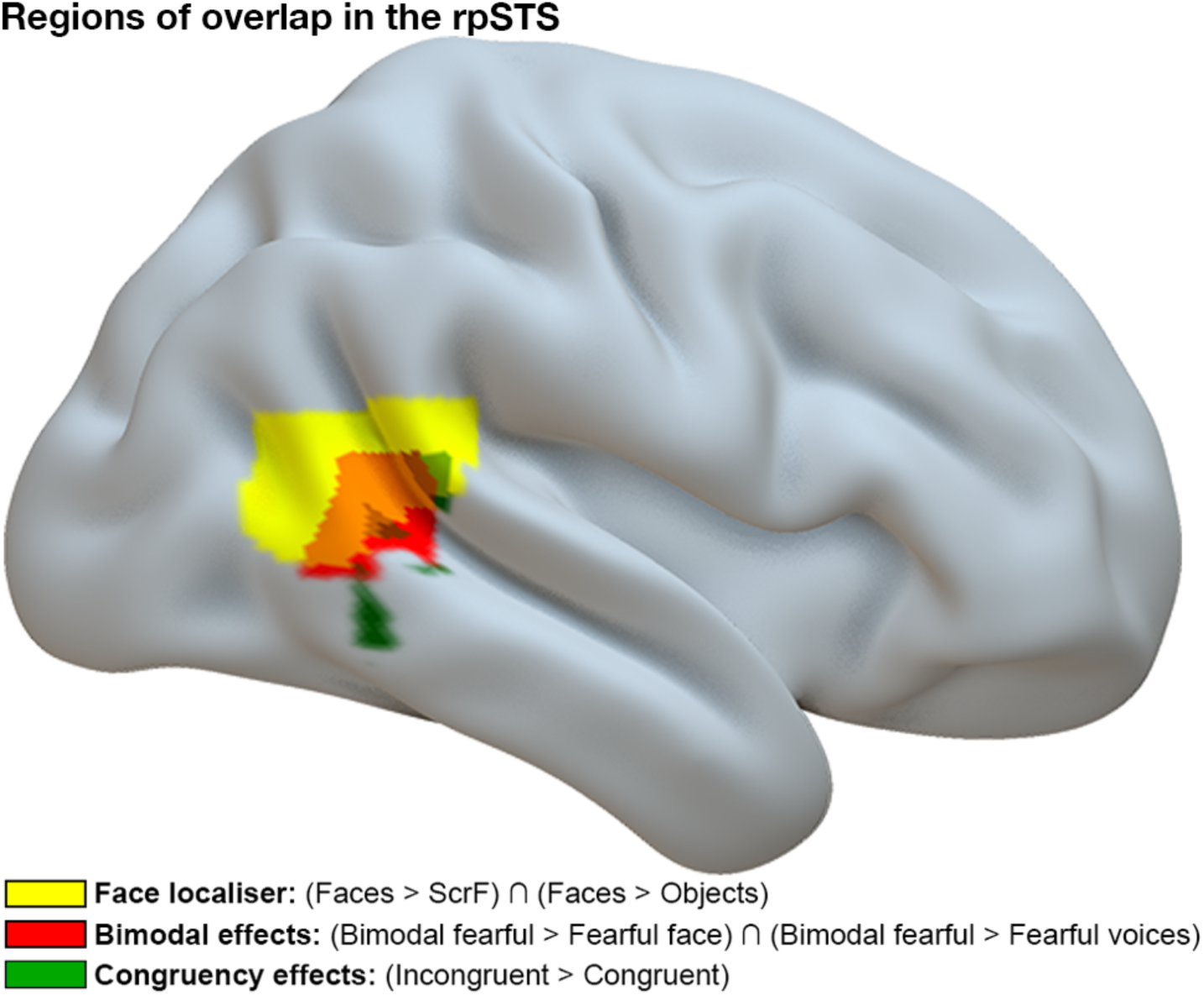
Regions of overlap in the rpSTS. Shown are the face-selective rpSTS (yellow), the region that responded to bimodal > unimodal fearful stimuli in the group analysis (red; see Figure 4), and the region in that responded more to incongruent > incongruent stimuli in the group analysis (green; Figure 7). Statistical maps thresholded at P < 0.05 FWE-corrected for multiple comparisons.

Next, we examined the response to congruent and incongruent stimuli in the face- and voice-selective regions of interest (Figure 7B). A 2×6 ANOVA (congruency, region) for the face-selective regions showed no effect of Congruency (F(1,23) = 1.75, P = 0.20), but significant effects of Region (F(5,115) = 6.05, P < 0.001) and an interaction between Congruency × Region (F(5,115) = 3.61, P < 0.005). Paired-samples t-tests showed this was due to a greater response to *incongruent > congruent* stimuli in the rpSTS (t(24) = 3.18, p < 0.05). There were no other significant differences (P’s > 0.20). A 2×5 ANOVA for the voice-selective regions also showed no effect of Congruency (F(1,23) = 2.29, P = 0.14), but significant effects of Region (F(4,92) = 7.04, P < 0.001) and an interaction between Congruency × Region (F(4,92) = 3.48, P < 0.05). This was due to a greater response to incongruent > congruent stimuli in the voice-selective right TVA (t(24) = 2.84, p < 0.05); there were no other significant differences (P’s > 0.20). Finally, a paired-samples t-test in the portion of rpSTS which showed a multimodal effect for fearful stimuli in the whole brain analysis, revealed a greater response to incongruent > congruent stimuli (t(24) = 2.11, p < 0.05).

### Dynamic Causal Modeling

We used Dynamic Causal modeling (DCM; (Friston *et al.* 2003) to investigate the cortical interactions between the face-selective right FFA (Figure 2), the voice-selective right TVA (Figure 3) and the portion of right posterior STS (rpSTS) showing a multisensory response (Figure 4) to understand the network dynamic of how emotional faces and voices are integrated. The FFA and TVA were chosen based on previous studies suggesting these regions are involved in individuating faces (Rotshtein et al. 2005) and voices (Belin *et al.* 2000; Belin and Zatorre 2003) respectively. Only right hemisphere regions were included in the analysis due to the multisensory response being observed in the rpSTS. The emotional expression condition was entered as a modulator of connectivity (B) between unisensory (FFA, TVA) and multisensory (rpSTS) regions, as well as a direct input to the rpSTS (C).

Since we were specifically interested in how the rpSTS integrates information from the FFA and TVA, and putative connections between FFA and TVA, we created 8 plausible models containing different ways in which these nodes may be intrinsically connected. We were specifically interested in whether reciprocal connections exist between the face-selective FFA and voice-selective TVA such has been previously suggested by studies on identity recognition (von Kriegstein and Giraud 2006; Blank *et al.* 2011; Blank et al. 2014), and how these regions are connected to the rpSTS – a region that this study and others (Ethofer et al. 2006; Kreifelts *et al.* 2009; Watson, Latinus, Noguchi, *et al.* 2014) have implicated as being involved in the integration of face and voice emotion information. Crucially, we examined the connections between the face-selective and voice-selective regions of interest identified in independent localiser scans, and the portion of rpSTS which showed a greater response to bimodal > unimodal faces and voices in the whole brain analysis (see Figure 4).

Each model was repeated for three alternative points at which emotional expression might influence the intrinsic connections: 1) directly modulating the response of the rpSTS; 2) modulating the connections between the unisensory FFA/TVA and the multisensory rpSTS; 3) directly modulating the response of the rpSTS as well as the connections between the unisensory FFA/TVA and the multisensory rpSTS. This resulted in a total of 24 models (Figure 9A).

**Figure 9.**
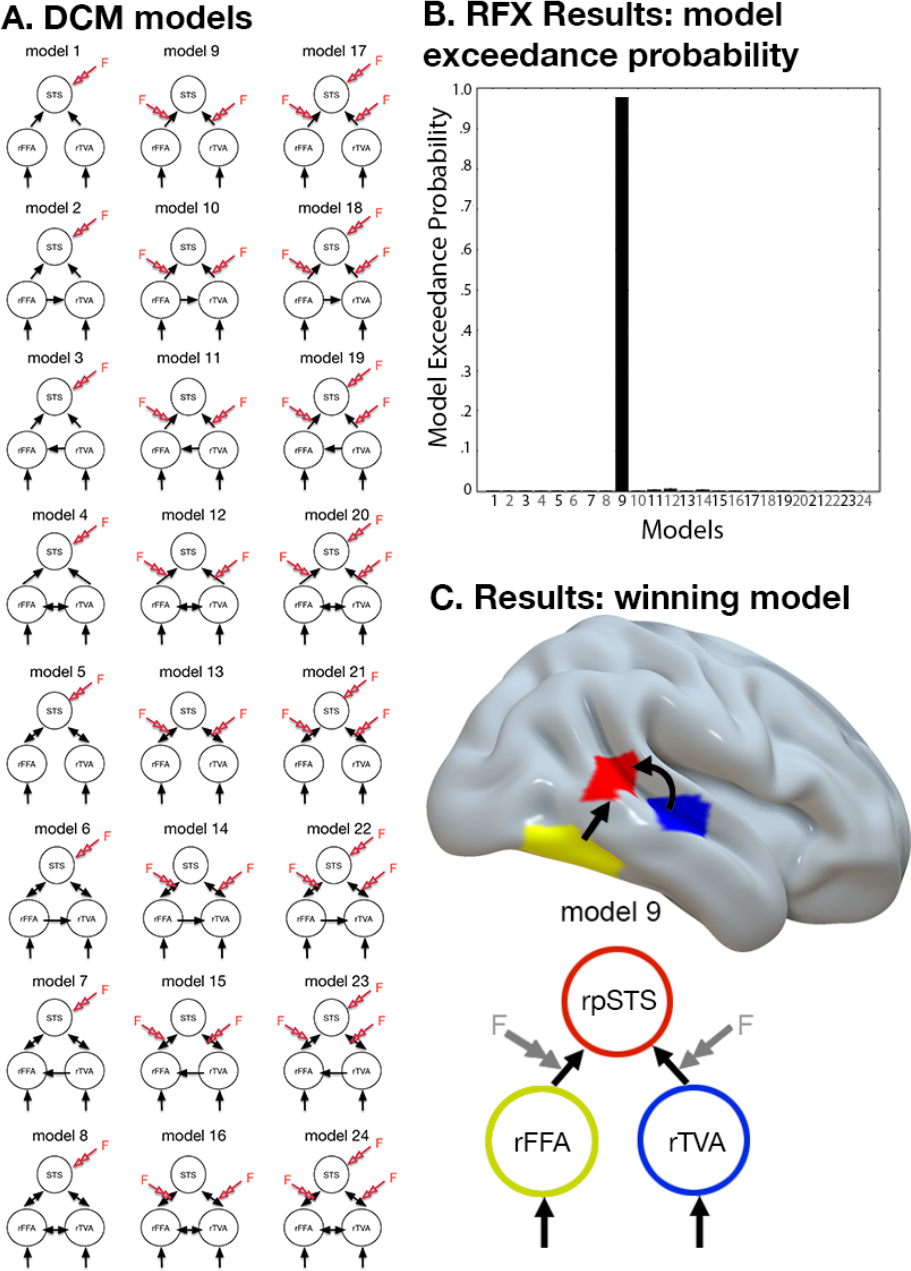
DCM analysis. A) Models entered into the DCM analysis. The model space was defined by variations of either feed-forward or feed-backward endogenous connections between the three regions (8 model types; rows), the type of modulation imposed onto the endogenous connections (columns) by expression. The resulting model space consisted of 24 models in total, which were entered into a RFX BMS analysis. B) Model exceedance probability obtained using RFX BMS across all participants, showed a strong preference for model 9. C) The winning model consisted of information entering the right FFA (yellow) and right TVA (blue), and significant feed-forward effective connectivity with the right pSTS (red). Emotional expression modulated the network response of the endogenous connections between the unimodal regions (right FFA and right TVA) and the multisensory right pSTS (as also shown in Figure 4).

Figure 9B shows the model exceedance probability obtained using RFX BMS across all participants for each of the 24 models in our models space. The highly winning model is model 9, with an exceedance probability of 0.9783 and expected posterior probability of 0.3151. The second highest exceedance probability was below 0.006. The winning model (Figure 9C) consisted of forward intrinsic connections from the FFA and TVA towards the rpSTS, with a modulatory effect of emotional expression of the face and/or voice on the network response of the endogenous connections between the unimodal regions (FFA and TVA) and the multisensory rpSTS (as shown in Figure 2).

## DISCUSSION

This study set out to specifically test at which stage of the face and voice processing hierarchy these signals are integrated, and how emotional expressions influences this integration. We also assessed whether crossmodal responses could be observed in face or voice selective regions (preferential response to face in voice selective region and reversely). Finally, we explored how information flows between face-selective, voice-selective and integrative regions.

Influential models of identity recognition propose that integration of information across the different sensory modalilites is performed at an independent stage where information from both modalities converge (Bruce and Young 1986; Burton *et al.* 1990). However, more recent animal (Ghazanfar *et al.* 2005; Perrodin *et al.* 2014, 2015) and human (Von Kriegstein *et al.* 2005; von Kriegstein and Giraud 2006; Blank *et al.* 2011; Joassin *et al.* 2011) studies suggested that multisensory integration of faces and voices may occur at earlier stages of processing, including in regions typically thought to be uniquely face or voice selective. For instance, in humans, studies have suggested that integration of face and voice identity information occurs via reciprocal connections between unisensory regions (for a review, see (Blank *et al.* 2014), with a portion of the fusiform cortex and the voice-responsive middle/anterior STS showing increased functional connectivity during a speaker recognition task as compared to a content-related task (Von Kriegstein *et al.* 2005), while the extent of the functional connectivity between these regions correlates with voice recognition abilities across subjects (Blank et al. 2015).

In our study, we found evidence that multisensory integration between face and voice signals of emotional expressions is selective to the right posterior STS (rpSTS) and arise after unimodal computation in unimodal face and voice selective regions. First, we observed a greater response to bimodal fearful stimuli as compared to unimodal stimuli in the whole-brain analysis in the rpSTS. This was further supported by a whole-brain analysis examining an interaction between emotion and modality, which revealed a single region within the right STS which responded significantly more to fear (bimodal > unimodal) than to neutral (bimodal > unimodal); this region partially overlapped with the region responding more to bimodal than to unimodal fearful stimuli. This region also activated differentially when faces and voices were presented in a congruent or incongruent fashion, overlapped with the face-selective rpSTS patch (as defined in an independent face localiser). Finally, we found that the multisensory rpSTS receives uni-directional information from the face-selective FFA, and voice-selective TVA, with emotional expression affecting the strength of the connections.

We found a significantly greater response to bimodal stimuli as compared to unimodal stimuli in the rpSTS, but only for fearful faces. This is in line with a previous study showing that the STS responds to emotion stimuli regardless of the sensory input (Peelen *et al.* 2010), and others showing a greater response in bilateral pSTS to bimodal as compared to unimodal faces and voices (Ethofer *et al.* 2006; Watson, Latinus, Noguchi, *et al.* 2014). However, the results from our study suggest that multisensory integration in the right STS may be specific for stimuli containing emotional expressions. This is the first study to show that multisensory integration of face and voice information in this region may be dependent upon the emotional content. Another possibility is that this region is specific for the integration of fearful stimuli. Indeed, a reduced response to fearful faces in the posterior STS is observed in patients with damage to the amygdala (Vuilleumier et al. 2004) suggesting that the STS may form a part of a distributed network for the automatic processing of fearful stimuli. Alternatively, this difference may be due to fearful stimuli being more salient than neutral stimuli, rather than the type of expression per se, and such responses may be modulated by top-down areas involved in attentional control (Desimone and Duncan 1995; Pessoa et al. 2002). This may partially explain the similar patterns observed for a greater response to bimodal than unimodal fearful stimuli observed across several face-selective regions.

While previous studies have also shown evidence that the integration of emotion face and voice stimuli occurs in the pSTS (Ethofer *et al.* 2006; Romanski 2007; Kreifelts *et al.* 2009; Watson, Latinus, Noguchi, *et al.* 2014; Hölig *et al.* 2017), other studies have suggested that MSI may occur via direct reciprocal connections between unimodal face and voice regions (von Kriegstein and Giraud 2006). One possible explanation for this discrepancy, is due to the fact that the former studies did not independently localize face and voice-selective regions, and test MSI within these regions. To test this, we conducted the first study which used independent localisers to identify face and voice-selective regions of interest, and examine the response in these regions to i) their preferred and non-preferred stimulus, ii) the response to bimodal stimuli, and iii) their response to congruent and incongruent stimuli.

First, we examined the response in ‘unimodal’ face and voice selective regions to their non-preferred stimulus. The FFA and OFA regions of the face selective network (see Figure 2) and the TVA, ATL, and aSTS regions of the voice selective network (see Figure 3) did not show any sign of crossmodal responses (faces in voice-selective region or voices in face-selective regions) or of multisensory integration (Figures 5-6). However, we observed significant responses to voices in bilateral pSTS as defined from the functional face localiser. Further, the face-selective rpSTS responded more to bimodal than unimodal stimuli, and responded more to incongruent than congruent stimuli.

Previous studies suggested that a portion of the fusiform gyrus plays a role in multisensory integration of faces and voices. For example, the fusiform shows a greater response to bimodal as compared to unimodal stimuli (Joassin *et al.* 2011) as well as an increased neural response following audio-visual learning of speaker recognition (von Kriegstein et al. 2008). However, in contrast to previous studies, we found no evidence of multisensory integration in this region – neither in the whole brain group analysis, nor in the face-selective FFA identified in an independent localiser. Interestingly, studies suggesting a role of the fusiform gyrus in multisensory integration of faces and voices manipulate the identity pairing of face and voice stimuli (von Kriegstein *et al.* 2008; Joassin *et al.* 2011), whilst the current study manipulated the emotion pairing. It is therefore possible that the FFA might play a multisensory role for identity discrimination while the rpSTS would integrate facial and vocal signals for emotional content. This is in line with models of face recognition proposing distinct pathways for processing identity and emotional expressions (Bruce and Young 1986; Burton et al. 1999; Haxby *et al.* 2000), as well as neuroimaging studies showing the FFA is sensitive to changes in identity but invariant to changes in emotional expressions (Winston et al. 2004), while the rpSTS is sensitive to emotional expression (Narumoto et al. 2001; Winston *et al.* 2004) but not to changes in identity (Davies-Thompson et al. 2009). However, other studies have shown significant interactions between identity and emotional expression processing; for example, the response in the FFA and rpSTS varies depending on whether participants are asked to attend to the identity or the emotional expression of the face stimuli (Gorno-Tempini et al. 2001; Ganel et al. 2005; Fox et al. 2009). Such task effect raises the interesting idea that the differences observed between our study showing multisensory integration of emotive face and voice stimuli in rpSTS, and other studies showing multisensory integration of identity face and voice information in the fusiform gyrus (i.e. (von Kriegstein *et al.* 2008; Joassin *et al.* 2011)), could be task-dependent constraints (emotion versus identity). A final consideration is that the use of an unrelated orthogonal task (gender discrimination) may have influenced the response with these regions. Early interaction between face and voice regions may crucially be under the influence of the task. For instance, studies have shown that the fusiform face area only responds to voices when participants performed a speaker identification task, but not when performing a verbal content task (Von Kriegstein *et al.* 2005). This again raises the possibility that task (i.e. emotion, identity, gender) may influence the loci of multisensory integration, and paves the way for future works aiming at directly comparing MSI of emotion versus identity within the same group of participants.

Neuropsychological studies have shown that individuals with face recognition deficits (prosopagnosia) typically remain able to identify individuals from voices (Liu et al. 2014; Liu et al. 2015), while individuals with voice recognition deficits (phonagnosia) have intact face recognition (Garrido et al. 2009). In line with these studies, with the exception of the rpSTS, we found no significant crossmodal responses in face-selective (OFA, FFA) or voice-selective (TVA, ATL, aSTS) regions; specifically, we observed no significant responses to voices in face-selective regions that, following damage, can result in prosopagnosia. Interestingly, damage to the rpSTS can result in deficits in facial expressions with intact facial identity as long as the expression remained unaltered (Fox et al. 2011), but can also leave vocal identity and vocal expression recognition intact (Jiahui et al. 2017). This suggests that while right rpSTS may be involved in multisensory integration of faces and voices, it is somewhat independent of unimodal voice processing.

We also examined the response to congruent versus incongruent bimodal stimuli. Behavioral studies have demonstrated that enhanced performance in emotion categorization tasks for congruent emotional stimuli, but impaired performance for incongruent stimuli (Massaro and Egan 1996; De Gelder and Vroomen 2000; Collignon *et al.* 2008). We reasoned that, if a region is involved in the integration of faces and voices then one would expect this region to be sensitive to congruency effects. We found that the rpSTS was indeed sensitive to congruency, both at the level of the region of interest analysis, as well as at the level of the whole-brain analysis, providing further support for the rpSTS to be involved in the multisensory integration of faces and voices of emotional stimuli. None of the other regions of the face or voice selective network showed sensitivity to the congruency of the stimuli. However, in addition to the rpSTS, several other regions responded more to incongruent than congruent stimuli, including the bilateral middle frontal gyrus, bilateral superior parietal cortex, and a portion of the left frontal pole (Figure 7). These regions are similar to those observed in previous studies suggesting a cingulate-fronto-parietal network involved in conflict monitoring (Carter et al. 1998; Botvinick et al. 2004; Weissman et al. 2004; Ochsner et al. 2009; Wittfoth et al. 2009; Müller et al. 2011). However, unlike the rpSTS, none of these regions responded significantly more to congruent bimodal than to unimodal stimuli for either fearful or neutral stimuli or showed face or voice selectivity in the independent localisers. Both of these methods have been used previously to identify regions of the brain involved in multisensory integration; however, the little overlap between regions responding more to bimodal than to unimodal stimuli, and regions showing congruency effects, suggest that these manipulations isolate distinct networks and processes (Morís Fernández et al. 2017). This is therefore the first study to show that, with the exception of the pSTS, face- and voice-selective regions show no evidence of MSI as measured by congruency effects – a finding which is in line with the observation that these regions also show no significant responses to their non-preferred stimulus (i.e. no response to voices in face-selective regions, no response to faces in voice-selective regions) in the localizer task.

Using DCM, we explored how face and voice signals flow between face-selective, voice-selective and heteromodal rpSTS regions, and how this information transfer is modulated by the emotional content of the stimuli. More specifically, we wanted to test if the integration of emotion across faces and voices relies on direct reciprocal interaction between FFA and TVA regions as previously suggested for the integration of identity information (von Kriegstein and Giraud 2006; Blank *et al.* 2011; Blank *et al.* 2014), or alternatively if face and voice signal is transferred and then integrated in a distinct convergence region in the pSTS (Ethofer *et al.* 2006; Kreifelts *et al.* 2009; Watson, Latinus, Noguchi, *et al.* 2014). We found strong evidence for the model receiving uni-directional information from the face-selective FFA and voice-selective TVA. Further, the winning model included fearful expressions affecting the strength of the connections between the unisensory regions (FFA, TVA) and the rpSTS. These results support the findings from a previous study showing increased functional connectivity between the FFA and pSTS for bimodal than unimodal face and voice stimuli (Kreifelts et al. 2007); however, we build upon these results in two ways: first, we found that the connections between these regions are unidirectional (FFA to STS, and TVA to STS), and secondly we found that the connections between these regions are significantly moderated by the presence of emotion stimuli as compared to neutral stimuli. These results provide support for a hierarchical architecture for the integration of emotional signals from faces and voices, with unisensory face and voice selective regions projecting to rpSTS where those information are integrated.

Finally, although previous studies looking at the anatomical connections between the various face-selective regions have suggested limited structural connectivity between the FFA and pSTS (Gschwind et al., 2012; Pyles et al., 2013), DCM is not a direct proxy for anatomical connectivity. Instead, it is a hypothesis driven functional connectivity technique testing which model, in a predefined plausible model space, best explains the data. In that sense, DCM does not tell us if direct anatomical connection exists between the nodes, but rather characterizes the causality between the activities of different brain areas and how information flows in the brain. One possibility, is that whilst direct anatomical connections between the face-selective FFA and face-selective pSTS do not exist, connections may be present between the FFA and the slightly more anterior and inferior regions of pSTS showing evidence of multisensory integration in the whole brain analysis. Alternatively, as DCM does not model the entire brain, but rather is constrained by the number of nodes entered into the models, areas beyond those entered into the models may also play a role in the multisensory integration of face and voice emotion information, such as regions specific to the type of emotion being presented (i.e. the amygdala). In our whole brain analysis, no other regions however showed evidence for multisensory integration of fearful stimuli.

In sum, we found support for multisensory integration of fearful face and voice in the rpSTS, as evidenced from: 1) a response to both faces and voices in the functional localiser scans; 2) a greater response to bimodal stimuli as compared to unimodal stimuli, which also overlaps with a region in the group analysis which showed a greater response to bimodal as compared to unimodal stimuli, as well as a significant interaction of emotion by modularity; 3) a face-voice congruency effect which overlaps with regions in the whole brain analysis showing bimodal integration. Using independent face and voice functional localizers we were able to show that these effects were only present in rpSTS and not in other face or voice selective regions. Finally, using DCM analysis, we found evidence that faces and voices are integrated in rpSTS using a bottom-up architecture, with the emotion content modulating the strength of these connections. Together, these findings promote a hierarchical model of integration of face and voice signals with a convergence zone in the rpSTS, and that such integration depends on the (emotional) salience of the stimuli, providing support for the prominent role the rpSTS plays in the multisensory integration of emotional stimuli.

## Acknowledgements

This study was supported in part by the Center for Mind and Brain Science (CIMeC), University of Trento and by a European Research Council starting grant attributed to OC (MADVIS: ERC-StG 337573). We thank anonymous reviewers for their helpful and insightful comments on our original manuscript.

## Notes

Conflict of interest: The authors declare no competing financial interests.

